# Facial photographs as proxies for inflammatory aging

**DOI:** 10.1101/2025.10.02.680162

**Authors:** Minja Belic, Kevin Schneider, David Furman

## Abstract

Systemic chronic inflammation is a major determinant of aging and disease risk, yet current biomarkers such as the Inflammatory Age (iAge) clock and other proteomic assays remain costly, invasive, and poorly scalable. The skin, as both a visible marker and contributor to age-related inflammation (“inflammaging”), offers a potential non-invasive window into this health metric. Here, we developed Healthy Selfie, a digital predictor of iAge, from simple 2D frontal face photographs. We leveraged data from the Edifice Health Pre-Market trial, a clinical study of 750 participants aged 20 to 90 years, using a subset with available iAge measurements paired to facial images, demographic, clinical, and functional data. Facial embeddings from a pretrained deep learning image model were combined with chronological age, sex and other easily obtainable metadata, and mapped to blood-derived iAge using regression within a leave-one-out cross-validation framework. Photo-predicted iAge values were significantly correlated with blood iAge values (r = 0.536) and were used to identify increased iAge acceleration (accuracy 66.39%, sensitivity 65.61%, specificity 67.24%). Validation on an external dataset containing ∼100,000 images showed no significant demographic bias across ethnicities, including Asian, Black, Indian, Middle Eastern, and Latino populations. Beyond iAge, we also showed that facial features could predict blood levels of individual inflammaging proteins (CXCL9, CXCL1, CCL11, TNFSF10, and IFNG). Our findings suggest that ordinary facial photos can provide information on blood inflammation and can be used as a scalable, ultra low-cost tool for assessing biological aging and advancing precision health.

## Introduction

Biological aging reflects the cumulative decline of organ systems and is a key determinant of disease risk and lifespan. Current methods to quantify biological age, such as DNA methylation, proteomic, metabolomic, transcriptomic and blood biochemistry clocks, have demonstrated predictive power, but remain expensive, relatively invasive, and limited in scalability. Recent advances in computer vision methods have achieved remarkable success in estimating chronological age from facial images (1). Extending this idea, deep learning models trained on 2D (2) and 3D facial photographs (3), as well as thermal images (4) have demonstrated that facial features can also capture biological aging processes and disease risk, with predicted biological ages correlating with metabolic parameters, cancer survival, and lifestyle factors. These studies suggest that the face encodes information about systemic aging beyond visible appearance, raising the possibility that other hallmarks of biological aging may likewise be reflected in facial morphology and skin. Unlike biospecimen-based assays, digital image acquisition is non-invasive, accessible, and readily scalable to billions of smartphone users worldwide. This combination offers a unique opportunity to democratize biological age assessment and provide actionable feedback for precision health and healthy longevity. However, despite these advances, the application of facial imaging to biological aging remains underexplored and has mainly focused on general measures of biological age, rather than specific aspects of aging.

Chronic low-grade inflammation, also known as ‘inflammaging’, is a major hallmark of aging, one that integrates and influences many other hallmarks of the aging process (5,6). Inflammaging arises from dysregulated innate immune signaling and increased systemic cytokine production, leading to persistent immune activation and tissue stress (7). Clinically, inflammaging has profound consequences: it underlies increased susceptibility to infectious disease, impaired vaccine responses, higher incidence of autoimmune conditions, reduced immune surveillance against cancer, and elevated risk of nearly all major non-communicable diseases in later life, including cardiovascular disease, type 2 diabetes, neurodegeneration, and frailty (8). Therefore, assessing systemic inflammation is an essential step toward healthy aging and a prerequisite for the development of preventative and personalized longevity medicine.

In skin, age-related changes in keratinocytes, fibroblasts, and melanocytes diminish tissue homeostasis and amplify oxidative stress, triggering the release of cytokines, chemokines, and lipid mediators that sustain chronic inflammation. Declining antioxidant levels decrease defenses and heighten vulnerability to environmental stressors such as UV radiation, creating a self-perpetuating cycle of reactive oxygen species production, protein damage, and diminished repair. Endocrine and neuroendocrine alterations, together with age-related shifts in the skin microbiome, compound this effect. Notably, cutaneous dysbiosis has been linked to frailty, inflammatory skin disease, and systemic immune dysfunction, while skin–gut axis interactions suggest that local inflammation can propagate to distant organs (9,10). Environmental exposures accelerate extracellular matrix degradation (11) and proteostasis loss, reinforcing both local and systemic inflammation. These processes indicate that skin aging is mechanistically intertwined with systemic aging, and highlight the skin’s dual role as both a visible biomarker of inflammaging and a biological driver of inflammatory burden, underscoring its potential as a non-invasive window into the state of systemic health (12).

Herein, we develop Healthy Selfie, a multimodal artificial intelligence pipeline that integrates convolutional neural networks applied to facial images with demographic and clinical metadata to predict inflammatory aging. The system extracts latent feature embeddings from facial morphometrics and aligns them with proteomic inflammatory signatures of aging based on iAge (13). By leveraging supervised deep learning and regularized regression, we establish a computational link between externally visible phenotypes and underlying molecular immune biology. In addition, we assess demographic robustness on an external dataset. This digital diagnostic platform enables scalable, non-invasive, and low-cost estimation of inflammaging, offering a practical alternative to conventional biomarker assays and a pathway toward precision longevity medicine.

## Methods

### Study Population

Data were obtained from the Pre-Market Trial (PMT) iAge® intervention trial conducted by Edifice Health (clinicaltrials.gov ID: NCT04983017). This decentralized, double-blind, randomized, placebo-controlled study enrolled 750 ambulatory adults without serious illnesses to characterize iAge in a real-world wellness population. Participants underwent assessments including cytokine panels, physical function tests, blood pressure, hemoglobin A1C, microbiome profiling, bulk RNA sequencing, functional intrinsic capacity domain measures, standard clinical parameters, and multiple patient-reported outcomes.

For the present analysis, we focused on individual baseline measurements prior to any intervention, and we considered a subset of 363 participants who met the following criteria: (i) availability of baseline iAge scores based on the measured top five components of iAge (CCL11, Interferon Gamma, CXCL9, TRAIL and CXCL1) (ii) a frontal facial photograph captured using an iPad that was suitable for computer vision analysis, and (iii) basic metadata, including chronological age at test time, sex, waist and hip circumference, weight, height, and performance on the timed up-and-go test. Descriptive cohort characteristics are provided in Table 1.

**Table 1.** Study cohort descriptive statistics.

| Variable | Female (N = 220; 60.6%) |  | Male (N = 143; 39.4%) |  |
| --- | --- | --- | --- | --- |
| | Mean ± SD | Range | Mean ± SD | Range |
| Age at test [years] | 56.74 ± 13.28 | 21 – 86 | 57.47 ± 13.08 | 26 – 86 |
| BMI | 23.80 ± 4.82 | 15.16 – 51.50 | 27.64 ± 5.80 | 19.95 – 75.03 |
| Height [m] | 1.65 ± 0.06 | 1.48 – 1.88 | 1.78 ± 0.08 | 1.53 – 1.95 |
| Weight [kg] | 65.02 ± 13.10 | 34.80 – 140.20 | 87.60 ± 20.59 | 46.70 – 256.80 |
| Hip circumference [cm] | 98.24 ± 9.65 | 75.50 – 132.00 | 103.39 ± 8.92 | 82.50 – 169.00 |
| iAge | 44.82 ± 13.14 | 16.43 – 89.35 | 40.87 ± 13.76 | 6.78 – 77.91 |
| Timed up and go [s] | 9.18 ± 1.63 | 5.09 – 14.00 | 9.29 ± 2.27 | 5.21 – 29.02 |
| Waist circumference [cm] | 82.56 ± 12.55 | 59.50 – 121.00 | 96.38 ± 11.31 | 71.75 – 169.00 |
| Waist-to-hip ratio | 0.84 ± 0.07 | 0.70 – 1.07 | 0.93 ± 0.06 | 0.78 – 1.10 |

### Image Processing and Feature Extraction

Images captured on iPad devices were converted from .HEIC to .png format on the command line using ImageMagick. Facial regions of interest were detected using the RetinaFace detector (14), aligned, and processed with the *buffalo_l* model set of the InsightFace Python package for face analysis, which employs the ArcFace deep learning architecture (15). Image embeddings were generated and concatenated with corresponding anthropometric metadata. This concatenated feature vector served as input for downstream modelling to estimate iAge (13) values, derived from blood protein data.

### Calculation of blood-derived iAge

Top five analytes from Sayed *et. al*. (13) were used to derive iAge. Luminex protein assays were run on samples from this cohort and protein standards were used to bridge each plate to the first plate. The fifth and ninety-fifth quantile for each analyte were used to scale the mean fluorescent intensities (MFI) for these analytes to those from Sayed *et. al*. and the linear model was used to calculate iAge.

### Feature Selection and Model Training

Feature selection was performed in a leave-one-out cross-validation (LOOCV) loop by retaining only those image and metadata features that exhibited a Pearson correlation coefficient greater than 0.1 with iAge in every training split. Retained features were scaled and used for training and testing. Multiple regression algorithms - including lasso, ridge, elastic-net, support vector machines, random forest regressors, multilayer perceptron regressors, and XGBoost - were tuned within each LOOCV iteration using nested cross-validation. Model performance was assessed using root-mean-square error (RMSE), coefficient of determination (R^2^), and Pearson correlation coefficient (r) between predicted and observed iAge values.

### Inflammatory Age Acceleration

Inflammatory age acceleration (iAgeAccel) was defined to quantify the deviation of an individual’s iAge from that expected based on their chronological age. Specifically, measured iAge values were regressed onto chronological age within the study cohort, and the residuals from this linear regression were taken as continuous measures of iAgeAccel. Positive residuals corresponded to individuals whose inflammatory aging exceeded that predicted by their chronological age (accelerated aging), whereas negative residuals reflected individuals with lower-than-expected inflammatory aging (decelerated aging).

For categorical comparisons, continuous residuals were dichotomized into two outcome groups: accelerated (residual > 0) and decelerated (residual < 0). This discretization enabled evaluation of classification tasks in addition to regression-based analyses. Both measured iAge (derived from blood proteomic profiles) and photo-inferred iAge (derived from facial embeddings and metadata) were used to compute residuals, allowing for direct comparison of the two modalities in estimating iAgeAccel.

Classification performance between measured and photo-inferred categories was assessed using standard diagnostic metrics, including accuracy (overall proportion of correctly classified individuals), sensitivity (true positive rate for identifying accelerated aging), and specificity (true negative rate for identifying decelerated aging). Confusion matrices were generated for each comparison to visualize classification outcomes and error patterns.

### Protein-Specific Analysis

Given that the original iAge model identified CXCL9 (Mig), TNFSF10 (TRAIL), IFNG, CCL11 (Eotaxin-1), and CXCL1 as the strongest contributors, these proteins were also analyzed separately to evaluate the feasibility of inferring their abundance from facial images, testing both sex-specific and sex-agnostic models. We calculated the Pearson coefficient of correlation between the photo-predicted and blood measured protein levels, ensuring generalizability through leave-one-out cross-validation.

### Demographic bias estimation

To assess demographic robustness of image-based iAge predictions, we evaluated performance on the FairFace dataset (16), which contains ∼100,000 images balanced for sex, race, and age. To refine age labels (provided in decades), we used the mean of each decade, averaged with age predictions from the MiVOLO visual transformer model for age and gender imputation (17). The unavailable metadata such as height and weight were sampled randomly from a normal distribution, based on parameters from literature (18,19). Group differences in predicted iAge and iAge acceleration were tested using Student’s t-test for gender and one-way ANOVA with Tukey post-hoc tests for race. Effect sizes were calculated using Cohen’s *d* and η^2^. Normality was assessed using the Shapiro–Wilk test and visual inspection of Q–Q plots.

All analyses in this study were conducted in R and Python.

## Results

### Prediction of blood-derived inflammatory age from 2D facial photos

We evaluated whether facial photographs could predict iAge using features derived from pretrained image embeddings, with or without simple metadata. Among several regression algorithms tested - lasso, ridge, elastic net, SVM, random forest, multilayer perceptron, and XGBoost (Figure 1) - elastic net (20) consistently performed best. Using photo-derived features only, the model achieved r = 0.469 and RMSE = 11.97 between predicted and target values. Including metadata as predictors improved model performance (r = 0.536, RMSE = 11.42), with 40% of estimates within a 5-year absolute error. Substituting chronological age with MiVOLO age estimates yielded similar results for iAge (r = 0.523, RMSE = 11.53). MiVOLO age estimates and true chronological age correlated strongly in our dataset with r = 0.93 (p = 7.8e-164) and mean absolute error (MAE) of 4.26 years, suggesting that chronological age can be substituted with a photo-derived estimate with no significant loss of performance (Figure 2).

**Figure 1.**
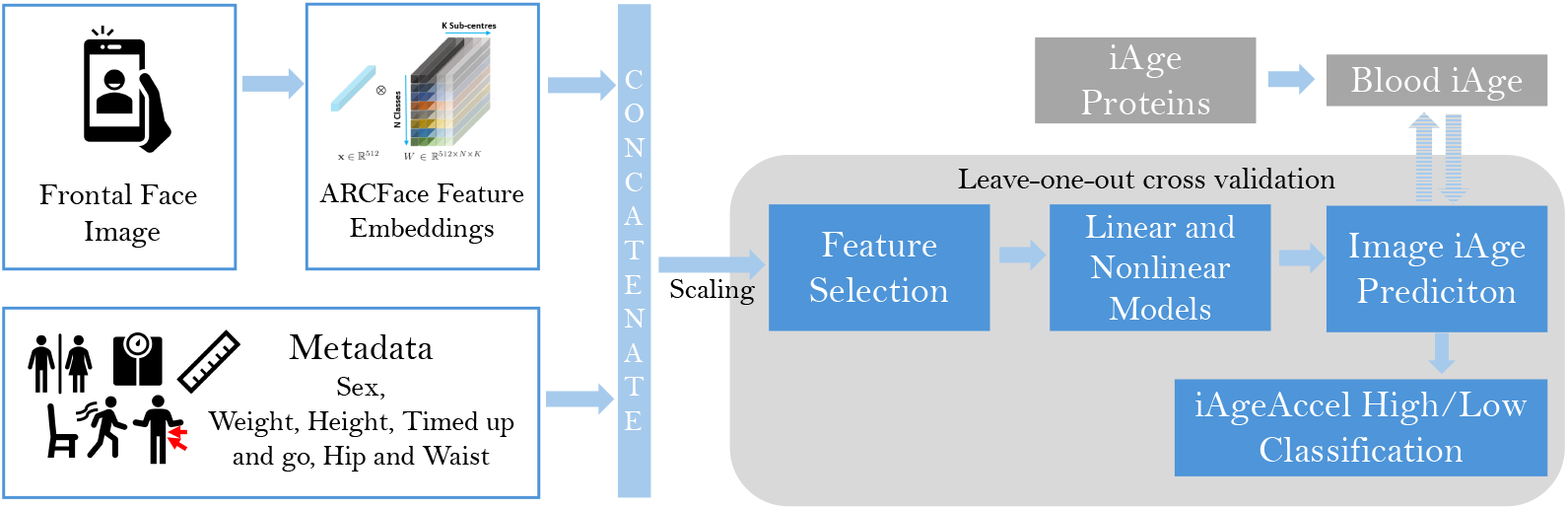
Study design. Frontal face images taken with any type of digital camera are fed into the ARCFace model [image adapted from (15)] and embeddings are concatenated with metadata. Following feature selection, different models are used to predict iAge, which is derived from five iAge proteins. Finally, iAge acceleration (iAgeAccel) is determined in reference to chronological age.

**Figure 2.**
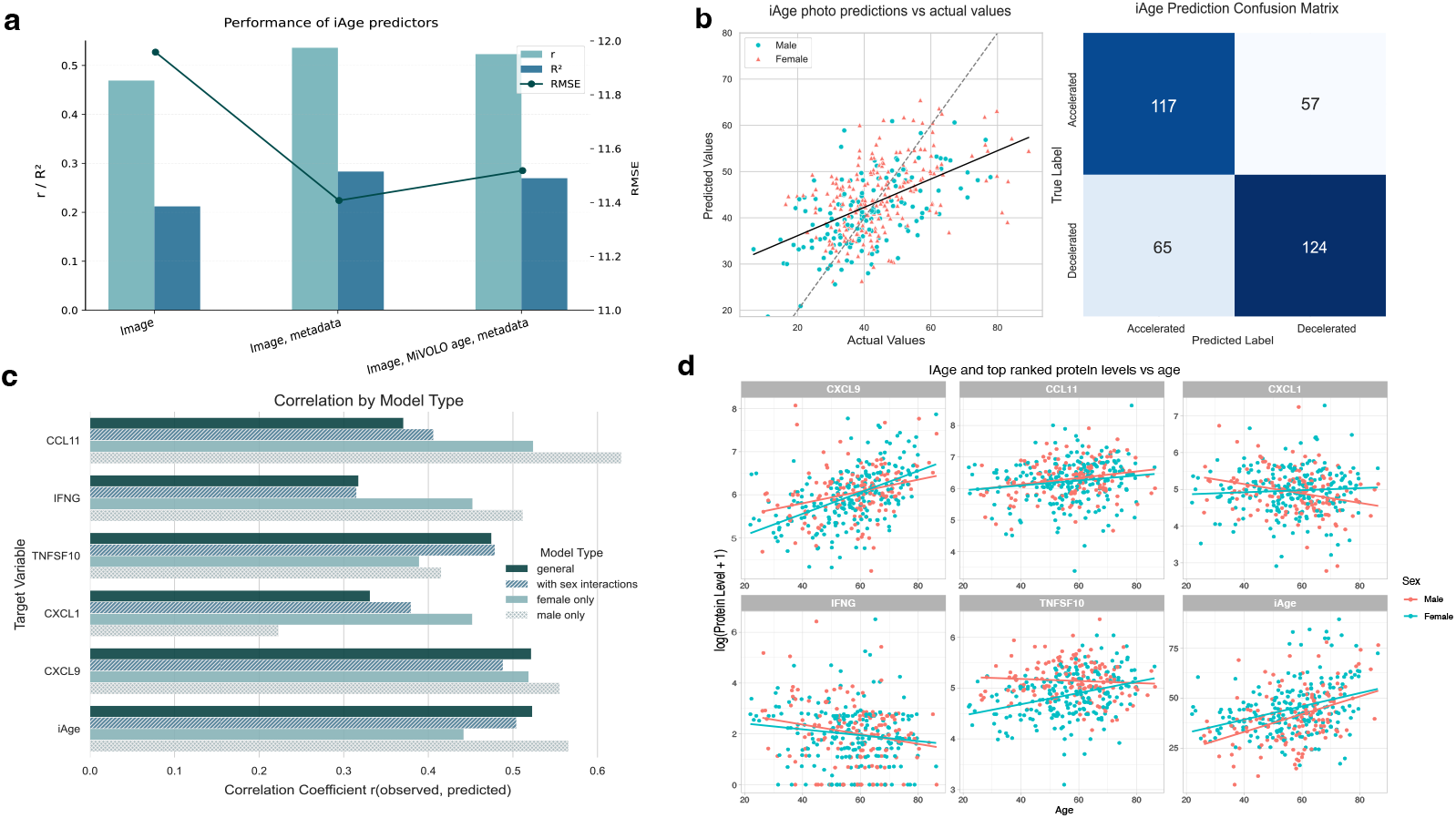
Performance metrics for the Image iAge models. a) Comparison of iAge model performance for different inputs were tested. Correlation, R^2^ and RMSE values are shown for prediction of iAge using only images, images and metadata, and images, metadata and age estimated by the MiVOLO model b) The observed and predicted values of iAge for each individual from the model using images and metadata are shown on the left, and predictions of iAgeAccel as a binary outcome are shown on the right c) The performance of model variations for five top ranked inflammatory proteins expressed as Pearson correlation between image-inferred and blood-measured values. d) iAge and top five proteins contributing most to iAge prediction, plotted against chronological age.

Thirty-two features from the original 512 InsightFace embeddings were retained in the final model using photo embeddings and metadata. Alternative embeddings (using FaceNet (21), VGGFace (22) and Derm Foundation (23)) and their combinations did not improve performance. For example, features extracted by Derm Foundation led to a modest correlation between predicted and measured iAge of r = 0.379, p = 7.4e-14. Using PCA-transformed features improved the model for Derm Foundation (r = 0.507, p = 3.8e-25) but not for InsightFace. This difference may reflect the larger dimensionality of Derm Foundation features (6144 vs. 512 for InsightFace), where dimensionality reduction is more beneficial.

When categorizing iAgeAccel into accelerated and decelerated categories, classification accuracy was 66.39%, sensitivity 65.61%, and specificity 67.24%. Figure 2b shows the corresponding confusion matrix for binary classifications.

### Prediction of iAge-specific circulating proteins from facial photos

Facial images and metadata were informative for predicting several key inflammaging proteins, with the strongest performance for CXCL9 (r = 0.522, p = 4.3e-27) and TNFSF10 (r = 0.475, p = 8.1e-22). Correlations were not as strong for CCL11 (r = 0.371, p = 2.9e-13) and were the weakest for IFNG (r = 0.318, p = 5.9e-10) and CXCL1 (r = 0.325, 2.1e-10). Inspection of sex-stratified relationship between protein levels and chronological age, where certain proteins exhibit differing slopes with age for males and females, as shown in Figure 2d, prompted us to add sex interaction terms to the model features, and also test models built only on the male or female populations (Figure 2c).

Adding sex interaction terms for all features in the model slightly improved the baseline performance for CCL11, TNFSF10 and CXCL1, but not for CXCL9 and IFNG. The best performance was achieved using models fitted only on the male subpopulation for CCL11 (r = 0.628, p = 6.5e-17), CXCL9 (r = 0.555, p = 6.1e-13) and IFNG (r = 0.512, p = 6.5e-11), while female-only model yielded the best results for CXCL1 (r = 0.452, p = 1.7e-12), and was less correlated in males (r =0.223, p = 0.007). The best model choice for TNFSF10 was the one built on the whole dataset adding sex interaction terms to the input features (r = 0.479, p = 3.4e-22). Estimation of the composite iAge score was best when trained only on males (r = 0.566, p = 1.8e-13), and worst when trained only on females (r = 0.441, p =6.2e-12), which was worse than the baseline model. Adding sex-interaction terms (r = 0.504, 8.2e-25) did not improve from the baseline model.

### Assessment of demographic bias of 2D photo-predicted inflammatory age

The Q–Q plot of iAge predictions on the FairFace dataset indicated that their distribution was approximately normal, with only minor deviations at the tails (Figure 6), although the Shapiro–Wilk test detected significant deviations from normality, which is unsurprising given the large sample size.

Mean predicted iAge was similar across demographic groups, with a median iAge around 37 (Figure 3b). Across gender, predicted iAge values differed slightly between males and females, but the effect size was negligible, with Cohen’s *d* = 0.086 (95% CI [0.073, 0.099]). When examining race, the omnibus ANOVA indicated statistically significant differences across groups (F(7, ∼97,000) ≈ 127, *p* < 1e-150). However, the corresponding effect size was very small (η^2^ = 0.0077), indicating that less than 1% of the variance in iAge predictions could be explained by race. Post-hoc pairwise comparisons using Tukey’s HSD test revealed that many racial group contrasts reached statistical significance. Nevertheless, the absolute mean differences were uniformly small, on the order of 0.2 to 1.6 years, with the greatest difference between the Black (36.3 ± 6.4) and Indian (37.9 ± 6.4) groups. Comparisons among Middle Eastern, Southeast Asian, and White groups were not significant.

**Figure 3.**
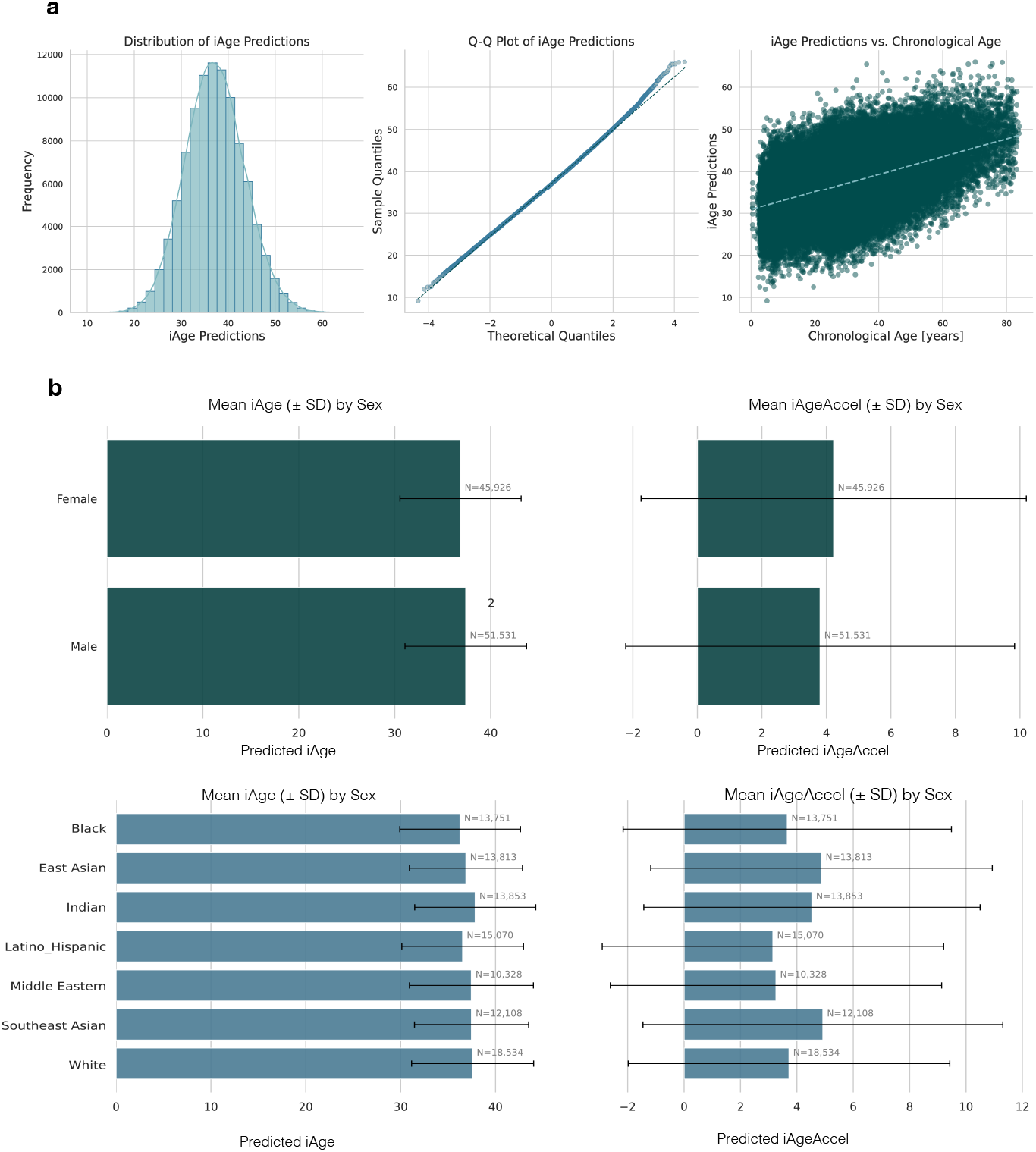
Predictions of iAge on the FairFace dataset. a) Histogram of predicted iAge values (left), Q-Q plot (middle) and iAge predictions against chronological age (right) b) Mean predicted iAge (left) and iAge acceleration (right) by demographic group, split by sex on top, and race on the bottom

ANOVA for iAgeAccel across races showed significant differences (F(7, ∼97,000) ≈ 207, p < 1e-263), with a small effect size (η^2^ = 0.0126), where race accounted for 1.26% of variance. The Tukey test found significant differences between many ethnic pairs, except for (Black, White), (Southeast Asian, East Asian) and (Latino Hispanic, Middle Eastern) groups, though the differences in means were relatively small, with the largest difference of 1.77 years between Latino Hispanic (3.1 ± 6.0) and Southeast Asian (4.9 ± 6.4) groups. iAgeAccel was marginally lower for males than females on this dataset (male 3.8 ± 6.0, female 4.2 ± 6.0, Cohen’s *d* = -0.068). The overall iAgeAccel mean was shifted upwards by 4 years on this test dataset, likely reflecting a shifted distribution of images compared to the training set, as well as the absence of true metadata values. Model application in new settings may thus benefit from recalibration.

## Discussion

This study provides proof of concept that facial images can be used to predict iAge, supporting the indications that the skin is not only a visible marker of systemic aging but also a biological contributor to inflammatory burden. By linking facial morphology with proteomic signatures of inflammation, we demonstrate the feasibility of a scalable, non-invasive, and ultra–low-cost approach to assessing biological age using a selfie and basic anthropometric information. Such digital tools could complement existing biomarker assays, which remain expensive and invasive, and as a rapid healthy aging screen, the Healthy Selfie could help democratize biological age measurement across large populations.

We used a dataset containing facial images paired with inflammatory protein profiles to fit a machine learning pipeline predicting iAge from facial images. iAge itself has been extensively validated: derived from the blood immunome of over 1000 individuals, it tracks with multimorbidity, frailty, immunosenescence, and cardiovascular aging, and is associated with exceptional longevity in centenarians (13). Adding simple anthropometric data, including waist and hip circumference and chronological age, improved the prediction of iAge compared to using images only. Substituting chronological age with its estimate from a deep learning age estimation algorithm marginally reduced performance; however this allowed us to test our model on out of sample data without precise chronological age attributes. Although the precision of exact iAge prediction from facial images was limited (RMSE >10 years), the ability to classify accelerated versus decelerated inflammatory aging was promising, with an accuracy of 66.39%, especially given that this level of performance was achieved using a relatively small dataset and off-the-shelf embeddings, illustrating the feasibility of facial biomarkers for systemic inflammation. Validation on the FairFace dataset containing ∼100,000 images showed comparable performance across gender and race. Although demographic effects were statistically detectable, they were small in magnitude, suggesting no practically significant demographic bias.

Aside from the composite iAge score, we also demonstrated the ability of facial features to serve as predictors of individual inflammaging proteins: CXCL9, TNFSF10, IFNG, CCL11 and CXCL1. Among the individual proteins, CXCL9 stands out as a particularly important biomarker of systemic aging and a major contributor to iAge. Previous work from the iAge study (13) has shown that CXCL9, largely produced by aged endothelial cells, predicts subclinical cardiovascular aging in otherwise healthy individuals, independently of conventional risk factors. That study also reviewed evidence linking elevated CXCL9 to multiple age-related conditions, including hypertension, left ventricular dysfunction, atopic dermatitis, and increased risk of falls in older adults. The fact that our model could predict CXCL9 levels directly from facial photographs is therefore especially significant, as it suggests that subtle facial features may capture endothelial and inflammatory changes linked to cardiovascular and frailty risk. Sex-specific modelling improved prediction for several proteins: male-only models yielded stronger performance for CCL11, CXCL9, and IFNG, while female-only models were optimal for CXCL1, and TNFSF10 was best captured by a combined model with sex-interaction terms. Similarly, iAge prediction was more accurate in a male-only than in female-only model. These patterns indicate that facial features may capture sex-dependent aspects of immune aging, with different proteins showing distinct trajectories in men and women.

Previous studies have used high-resolution 3D facial imaging to show that facial age acceleration associates with inflammatory pathways and metabolic health (3). While powerful, such approaches rely on specialized equipment, limiting scalability. By contrast, we demonstrate that standard 2D facial photographs, obtainable with common devices such as tablets or smartphones, can be used to approximate inflammatory age and its components.

Several limitations to this work should be acknowledged. The dataset used here was relatively small by computer vision standards, which limited our ability to fine-tune deep learning models and instead necessitated downstream modelling on fixed embedding spaces. Larger datasets would enable meaningful task specialization and likely improve predictive performance. Moreover, our tests across several embedding models yielded similar results, suggesting that without fine-tuning, current face analysis models capture limited information relevant to inflammatory age. In particular, the InsightFace embeddings used here were pretrained for identity recognition rather than biological aging; while they provide robust descriptors of facial morphology, they may not optimally encode features linked to inflammaging. Future work could therefore benefit from embeddings derived from models pretrained specifically for aging or other biological features related to inflammation and aging. Finally, it remains unclear whether the native 112×112 px resolution of many standard models is sufficient to capture subtle skin features linked to inflammation. Interestingly, even when using higher-resolution inputs with Derm Foundation (448x448 px), predictive performance did not improve, possibly because the larger feature space introduced too many degrees of freedom given our sample size.

Future work will include incorporating a broader range of metadata, including lifestyle, questionnaire, and clinical blood measurements, which may improve precision. Expanding image acquisition beyond frontal views to include side and scalp photographs could provide complementary information and increase the discriminative power of embeddings. Extending analyses to multiple visits per participant would enable longitudinal modelling of inflammatory age trajectories within individuals. Together, these enhancements would allow a more robust and dynamic understanding of how facial features encode systemic aging processes. The work presented here is an illustration of what health insights can be gleaned through the facial photos, and we believe the spectrum of possibilities is much larger. We plan to extend this approach to estimate a variety of organ-specific aging metrics and integrate them into a digital app, potentially complemented by other modalities such as voice analysis.

A major bottleneck for progress in this field is the absence of openly accessible datasets that combine facial photographs with detailed health outcomes. Advances in face recognition and age estimation over the past decade have been powered by large, diverse datasets such as WebFace, MegaFace, and CelebA. Comparable datasets linked to clinical information could drive similar progress in facial biomarker research for systemic diseases. Yet such data raise profound privacy and ethical challenges. Unlike most biochemical and physiological tests, facial photographs are inherently identifiable, and linking them with sensitive health information increases the risk of individual re-identification, limiting the feasibility of broad data sharing under current regulations.

Despite these challenges, recent initiatives suggest promising paths forward. For example, hybrid strategies such as the FAHR-Face (24) project demonstrate that models can be pre-trained on massive, publicly available image datasets in an unsupervised fashion, then fine-tuned on restricted clinical datasets containing health outcomes. This approach preserves some of the benefits of large-scale training while limiting direct exposure of sensitive health-linked facial data. Ultimately, progress in this area will depend on carefully balancing the scientific value of health-linked facial datasets with the imperative of protecting participant privacy.

In sum, our results provide an early demonstration that biological age prediction from facial images is feasible. While current performance is constrained by data availability and reliance on pretrained embedding models, the potential of facial imaging as a digital biomarker of systemic health is considerable. With larger and more diverse datasets, methodological refinements, and responsible approaches to privacy, digital image-based diagnostics could evolve into a transformative, ultra-low-cost alternative for aging assessment - scalable to global populations and aligned with the goals of precision health and healthy longevity.

